# PEG10-ORF1 programs trophoblast progenitor development for placental labyrinth formation

**DOI:** 10.1101/2025.10.08.681076

**Authors:** Hirosuke Shiura, Mayuko Fujii, Masatoshi Ooga, Sayaka Wakayama, Daiyu Ito, Takashi Kohda, Tomoko Kaneko-Ishino, Fumitoshi Ishino

**Affiliations:** Faculty of Life and Environmental Sciences, University of Yamanashi, Yamanashi 400-8510, Japan; Department of Animal Science and Biotechnology, School of Veterinary Medicine, Azabu University, Kanagawa 252-5201, Japan; Advanced Biotechnology Center, University of Yamanashi, Yamanashi, 400-8510, Japan; Faculty of Nursing, School of Medicine, Tokai University, Kanagawa 259-1193, Japan; Department of Epigenetics, Medical Research Institute (MRI), Tokyo Medical and Dental University (TMDU), Tokyo 113-8510, Japan

**Keywords:** *PEG10*, gene domestication (exaptation), placenta development, multiple placental roles, eutherian placenta evolution

## Abstract

The emergence of the placenta was a significant event in mammalian evolution, enabling therian mammals to give birth to live young through viviparity. A key genomic change associated with this process was likely the acquisition of the therian-specific paternally expressed 10 (*PEG10*) from a metavirus, as *Peg10*-null mutant mice exhibit early embryonic lethality due to the absence of both the spongiotrophoblast and labyrinth layers in the placenta. *PEG10* encodes a GAG-like open reading frame 1 (ORF1) and a POL-like ORF2, and produces two proteins, PEG10-ORF1 and PEG10-ORF1/2 fusion proteins, via the −1 frameshift mechanism conserved in retroviruses and metaviruses. Here, we demonstrate PEG10*-*ORF1 plays an essential and distinct role in labyrinth development. Mice lacking ORF1 but retaining the ORF1/2 protein initially developed normally in both embryos and placentas; however, later exhibited underdevelopment of the placental labyrinth’s complex microarchitecture, accompanied by mid-to late gestational growth retardation and lethality. These severe defects are most likely attributable to dysregulated development of labyrinth trophoblast precursor (LaTP) cells. Beyond its essential role in early placental formation, we previously demonstrated that the DSG protease domain within PEG10-ORF1/2—a metaviruse-derived feature—is critical for maintaining fetal capillaries during late gestation. Thus, *PEG10* fulfills multiple indispensable functions in placental formation and maintenance by exploiting a metavirus-derived multi-protein and -peptide production system, despite being a single gene. This study provides additional evidence underscoring the profound role of *PEG10* acquisition in therian placental evolution.

**Significance Statement:** The placenta is an essential organ for viviparous reproduction in therian mammals; therefore, its evolutionary emergence is of significant interest. One key event may have been the acquisition of *PEG10*, a metavirus (retrovirus-like LTR retrotransposon)– derived gene, by the common therian ancestor. We previously showed that *Peg10* is required for early placental development and that its protease domain is essential for late-gestational maintenance of fetal capillaries within the placental labyrinth. Here, we provide additional genetic evidence that *Peg10* is also indispensable for mid-gestational development of the placental labyrinthine microvasculature. Together, these findings illustrate how multiple, stage-specific functions of a single metavirus-derived gene, *PEG10*, coordinate placental formation and maintenance, highlighting the evolutionary importance of *PEG10* acquisition in therian placentation.

## Introduction

The placenta is an essential organ for viviparous reproduction in therian mammals. Marsupials possess a choriovitelline placenta (yolk sac placenta) and typically give birth to highly altricial young after a short gestation, relying on prolonged lactation to complete development (1, 2). In contrast, eutherians have a more sophisticated chorioallantoic placenta, which supports extended in utero development. Eutherian placental structures have diversified during evolution, encompassing hemochorial, endotheliochorial, and epitheliochorial types (3). Phylogenetic and uterine transcriptome analyses suggest that the ancestral eutherian placenta was hemochorial (4, 5). The mouse placenta, classified as hemochorial, consists of three major layers: the labyrinth, spongiotrophoblast, and trophoblast giant cell (TGC) layers. The labyrinth, formed by the fusion of the chorion and allantois, contains complex fetal vascular microstructures, where fetal capillaries are directly exposed to maternal blood. This structure is essential for efficient nutrient, waste, and gas (O_2_/CO_2_) exchange between the fetal and maternal circulations. The spongiotrophoblast layer structurally supports the labyrinth and directs maternal blood flow from the decidua into it, while the TGC layer separates the maternal decidua from the placenta. Importantly, the spongiotrophoblast and TGC layers also secrete placental hormones and cytokine-like proteins that are critical for maintaining gestation (6). Thus, the eutherian placenta performs a variety of essential functions through its sophisticated architecture. Given these functions, the evolutionary emergence of the therian placenta is of considerable interest. One key event may have been the acquisition of paternally expressed 10 (*PEG10*) by the common therian ancestor, as this gene plays a critical role in placental development (7, 8).

*PEG10* is a therian-specific gene domesticated from a metavirus—specifically, a retrovirus-like long terminal repeat (LTR) retrotransposon of the *Ty3/Gypsy* lineage (8– 11). It encodes two open reading frames (ORFs): ORF1, which contains a CCHC-type zinc-finger RNA-binding motif, and ORF2, which contains the canonical DSG catalytic motif of retroviral aspartyl proteases. These ORFs share ∼20–30% amino acid identity with the GAG and POL proteins of the *sushi-ichi* retrotransposon isolated from pufferfish (9). *PEG10* produces two proteins—PEG10-ORF1 and the PEG10-ORF1/2 fusion protein—the latter arising via programmed −1 ribosomal frameshifting, as observed in retroviruses and metaviruses (LTR retrotransposons) (12, 13). The domain architecture of *PEG10* is highly conserved across therian mammals (13), suggesting that both PEG10-ORF1 and PEG10-ORF1/2—as well as their hallmark motifs (the ORF1 CCHC zinc finger and the ORF2 DSG protease)—are under functional constraint and perform distinct roles. In addition, the ORF2-encoded protease produces multiple PEG10-derived self-cleavage products (13, 14), further increasing its biochemical complexity. Thus, although encoded by a single gene, *PEG10* appears to exert diverse molecular functions. To unravel this complexity, a stepwise genetics-based strategy is needed, such as generating a panel of *Peg10* mutant mice that target each of its highly conserved virus-derived features.

A complete loss of *Peg10*, eliminating both the ORF1 and ORF1/2 fusion proteins, results in early embryonic lethality in mice before 10.5 days post coitus (dpc), due to the absence of two major placental layers: the spongiotrophoblast and labyrinth (7). This finding demonstrates that *Peg10* is indispensable for trophoblast growth and differentiation during early placental development. By contrast, PEG10-protease mutant mice (PEG10-ASG mutants), in which the catalytic aspartic acid (D) of the DSG motif is substituted with alanine (A), exhibit a distinct phenotype. Their placentas appear normal until mid-gestation but, beginning at ∼12.5 dpc, exhibit both embryonic and placental growth retardation. More than 80% of these mutants die during perinatal development due to severe inflammation around the fetal vasculature in the labyrinth layer. These findings indicate that PEG10 protease activity is essential for maintaining fetal capillaries during late gestation, and that its loss disrupts feto-maternal circulation (14, 15).

In this study, we generated PEG10-ORF1-deficient (*Peg10*-ORF1 KO) mice by mutating the highly conserved “slippery sequence” (GGGAAAC), immediately upstream of the ORF1 stop codon, thereby abolishing −1 ribosomal frameshifting and the ORF1 protein production, while preserving ORF1/2 fusion protein production. The slippery sequence directs ribosomal pausing, backtracking by one nucleotide, and reinitiation in the ORF2 frame to produce the ORF1/2 fusion protein (Fig 1; *SI Appendix*, Fig. S1). To disrupt this mechanism, we replaced GGGAAAC with **T**GG**C**AAT and inserted a cytosine at the 3’ end (resulting in TGGCAATC), which allowed the ORF1/2 amino acid sequence to remain intact. Importantly, ORF1 KO mice exhibited frequent lethality beginning at mid-gestation, with mortality exceeding 80% by term. This outcome was attributable to disruption of the labyrinth’s complex microstructure, where fetal endothelial cells are normally enveloped by trophoblast layers. This phenotype is likely due to dysregulated development of labyrinth trophoblast precursor (LaTP) cells. Taken together, these *Peg10* mutant studies demonstrate that *PEG10* fulfills multiple essential, stage-specific roles in placental development by employing a metavirus-derived system for generating diverse proteins and peptides. These findings provide strong evidence that the acquisition of *PEG10* was a key event in the evolution of viviparity in mammals.

**Figure 1.**
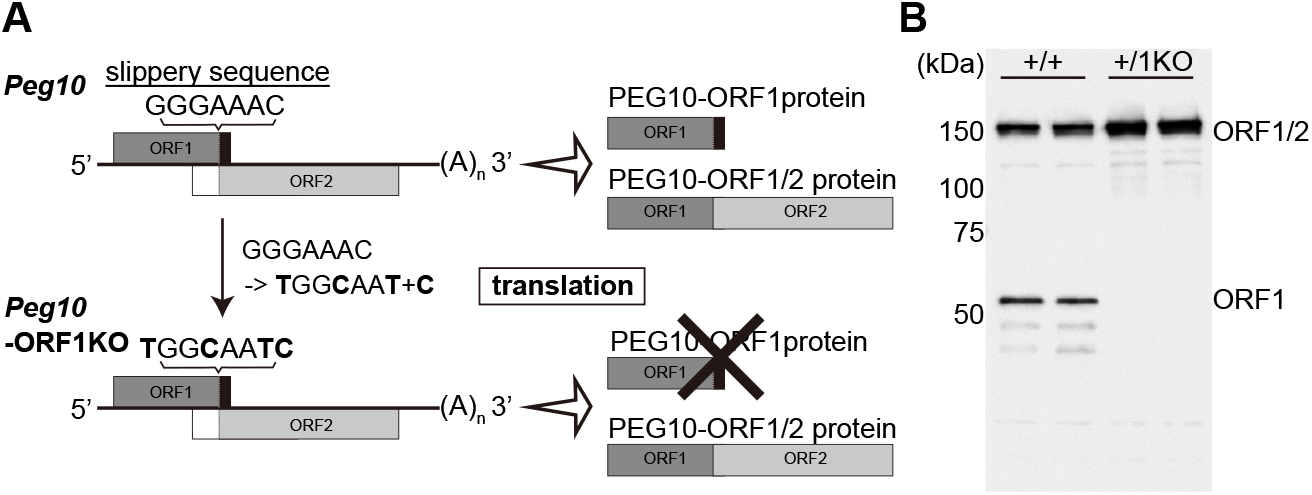
Generation of *Peg10*-ORF1KO mice. (A) Schematic representation of the *Peg10*-ORF1KO mutation. In order to disrupt the frameshift and thereby enable the production of solely the PEG10-ORF1/2 protein, the slippery sequence, GGGAAC, which is essential for the −1 frameshift, was replaced with TGGCAACC. (B) PEG10 protein expression in placenta at 10.5 dpc was confirmed by using an anti-PEG10-ORF1 antibody which detects both PEG10-ORF1 and PEG10-ORF1/2 proteins. The PEG10 ORF1 protein was undetectable in the +/1KO placentas.

## Results

### Generation of PEG10-ORF1 deficient (*Peg10*-ORF1KO) mice

The presence of two ORFs, a GAG-like ORF1 and POL-like ORF2, together with the consensus slippery sequence (GGGAAAC) required for −1 ribosomal frameshifting, is highly conserved in the *PEG10* gene of therian mammals (13), suggesting that both PEG10-ORF1 and PEG10-ORF1/2 proteins are functionally important. To assess the specific role of ORF1 in mammalian development, we generated PEG10-ORF1-deficient mice by mutating the conserved slippery sequence. This mutation abolished −1 frameshifting, thereby preventing ORF1 production while preserving translation of PEG10-ORF1/2 fusion proteins through the canonical reading frame without altering the amino acid sequence (16) (Fig. 1A; *SI Appendix*, Fig. S1C). Because multiple ribosomal frameshift outcomes have been reported (17), we specifically selected a mutation that reproduced the predominant ORF1/2 protein sequence observed in wild-type mice. Since *Peg10* is a paternally expressed gene, mice inheriting a paternal mutant allele (+/1KO; hereafter referred to as *Peg10*-ORF1KO or 1KO mice) were used for phenotypic analyses. First, we investigated whether *Peg10*-ORF1KO mice exhibited early embryonic lethality. In contrast to *Peg10* null KO mice, the mutants showed no gross embryonic or placental abnormalities at 10.5 dpc (*SI Appendix*, Fig. S2). Immunoblotting with an antibody that detects both PEG10-ORF1 and PEG10-ORF1/2 proteins confirmed the presence of both proteins in the wild-type (+/+) placentas. However, only the ORF1/2 protein was detected in *Peg10*-ORF1KO placentas, consistent with the designed mutation (Fig. 1B). These findings indicate that the loss of PEG10-ORF1 alone does not cause the early embryonic lethality associated with the severe placental dysplasia observed in *Peg10-*null KO mice.

### Effect of loss of PEG10-ORF1 protein on fetal growth and lethality

Loss of PEG10-ORF1 protein resulted in progressive fetal growth retardation and high lethality beginning in mid-gestation. From 12.5 dpc, *Peg10*-ORF1KO fetuses exhibited significantly reduced growth, with fetal weights decreased by 10–15% compared with wild-type littermates. The defect became increasingly pronounced, such that by 18.5 dpc, mutant fetuses reached only ∼60% of wild-type fetal weight, whereas placental growth was relatively preserved, showing only ∼10% reduction at all stages examined (Fig. 2A).

**Figure 2.**
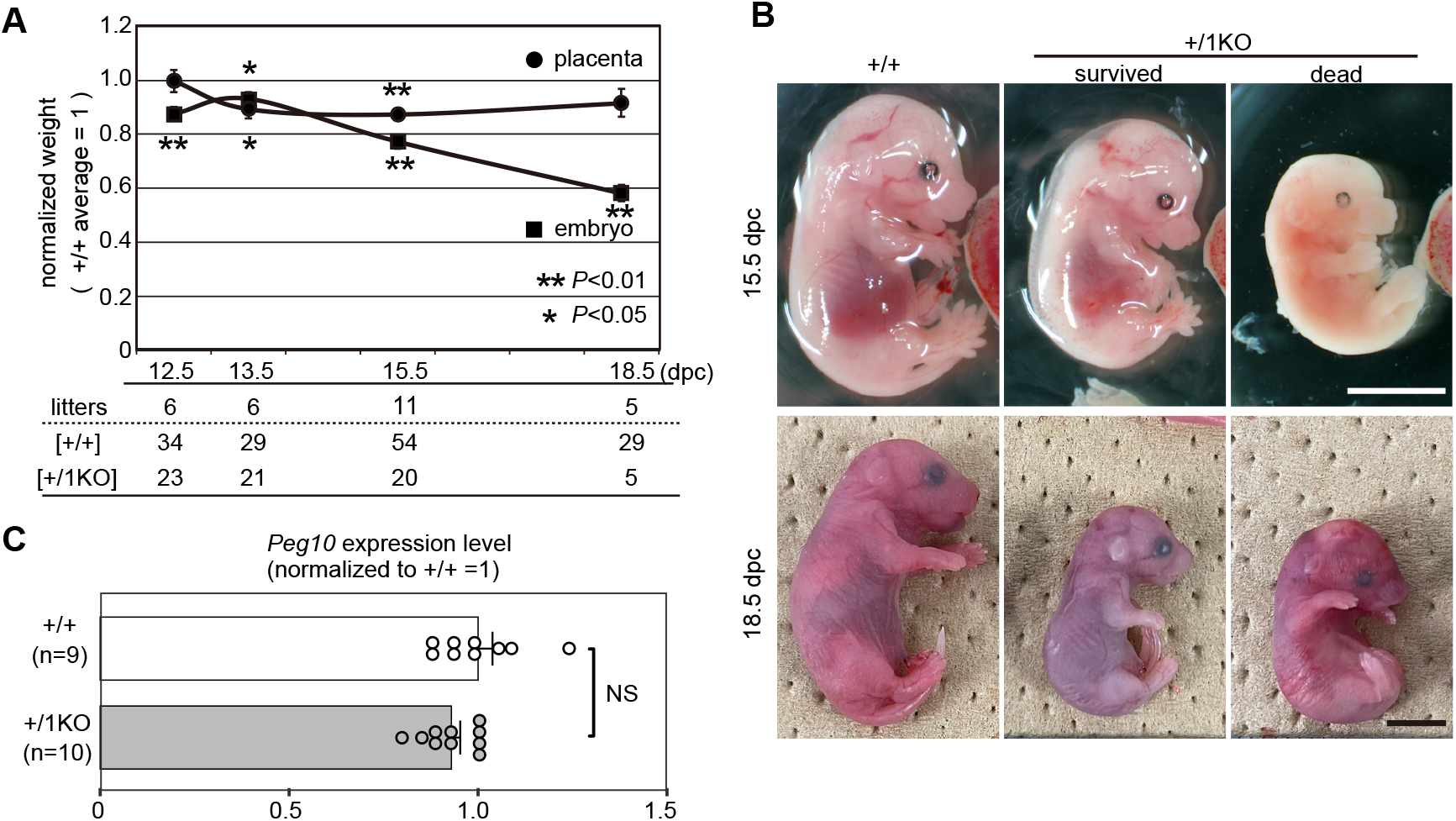
Severe growth defects and lethality observed in +/1KO mice. (A) Comparison between wildtype (+/+) and *Peg10*-ORF1KO (+/1KO) embryo and placental weights. Mean weights were calculated for each genotype within a given litter and a value of 1 represents the mean weight of the +/+ mice. The mean values ± s.e.m. of each genotype were plotted. Statistical significance was calculated using a two-tailed unpaired Student’s t-test with Welch’s correction. (*: P < 0.05, **: P < 0.01). (B) +/+ and +/1KO embryos at 15.5 (top) and 18.5 (bottom) dpc. Some of the +/1KO embryos recovered at these stage were already dead (rightmost). The scale bars indicate 5 mm. (C) *Peg10* expression level in 9.5 dpc placenta. The value of ‘1’ represents the mean *Peg10* expression level normalized to *Actb* in the +/+ placenta. In total 19 samples from two litters (+/+ : +/1KO = 9 : 10) were examined. The mean values of each genotype were plotted and each dot indicates the value obtained from each sample. Statistical significance was calculated using a two-tailed unpaired Student’s t-test with Welch’s correction. The error bars indicate the s.e.m. NS; Not significant.

The number of viable 1KO fetuses appeared reduced as early as 12.5–13.5 dpc (not statistically significant); however, lethality became evident around 15.5 dpc (Figs. 2A, B; Table 1). In these crosses, wild-type (+/+) females were mated with heterozygous 1KO/+ males (heterozygous carriers inheriting the mutant allele maternally), thereby excluding maternal effects and ensuring that the expected ratio of 1KO to wild-type fetuses was 1:1. However, the observed proportion of 1KO fetuses declined to ∼40% at 15.5 dpc and fell to < 20% by 18.5 dpc. Because *Peg10* expression levels remained unaltered by this mutation (Fig. 2C), these results demonstrate that the PEG10-ORF1 protein is essential for normal fetal development, and its absence causes severe growth retardation and frequent lethality from mid-gestation onward.

**Table 1.** The number of embryos or pups from +/+ dams crossed with 1KO/+ sires.

| | Litters | Observed<br>+/+ : +/-1KO | Expected<br>ratio<br>( 1 : 1 ) | $\chi^2$ value | P-value |
| --- | --- | --- | --- | --- | --- |
| **12.5 dpc** | 6 | 34 : 23<br>(1 : 0.68) | 28.5 : 28.5 | 2.12 | NS (0.15) |
| **13.5 dpc** | 6 | 29 : 21<br>(1 : 0.72) | 25 : 25 | 1.28 | NS (0.26) |
| **15.5 dpc** | 11 | **54 : 20**<br>**(1 : 0.37)** | 37 : 37 | 15.6 | **< 0.0001** |
| **18.5 dpc** | 5 | **29 : 5**<br>**(1 : 0.17)** | 17 : 17 | 16.9 | **< 0.0001** |

The Chi-square test was used to determine whether the difference between an observed and expected frequency distribution (mendelian ratio; 1:1) was statistically significant.
NS: Not significant. The numerical values in parentheses indicate the value when the +/+ is set to 1 in the "Observed" column.

### *Peg10*-ORF1 KO mice exhibit an aberrant morphology of the placental labyrinth

Mutant embryos appeared normal aside from growth deficiency; however, striking abnormalities were observed in their placentas. Immunofluorescence using an antibody specific to the extreme C-terminus of PEG10-ORF1, which detects only the ORF1 protein, confirmed its absence in mutant placentas (*SI Appendix*, Fig. S3), consistent with the immunoblotting results described above. At 13.5 dpc, whole-mount observation of the placenta from the fetal side showed that wild-type placentas formed a smooth, circular labyrinth layer, whereas mutant placentas exhibited jagged, irregular boundaries, and the blood within the layer appeared paler than that in wild type (Fig. 3A). These abnormalities were already evident in some mutants at 12.5 dpc and persisted as development progressed (*SI Appendix*, Fig. S4). Histological analysis further revealed morphological defects. Hematoxylin and eosin (HE) staining of 13.5 dpc sections showed comparatively large cavities in the mutant labyrinth (Fig. 3B). In addition, alkaline phosphatase (AP) staining, which marks trophoblast cells in the labyrinth (18), demonstrated prominent islet-like unstained regions in mutants (Fig. 3C). Together, these findings suggest that labyrinth development in *Peg10*-ORF1KO placentas is impaired or delayed, potentially leading to reduced fetal and/or maternal circulation within the placenta.

**Figure 3.**
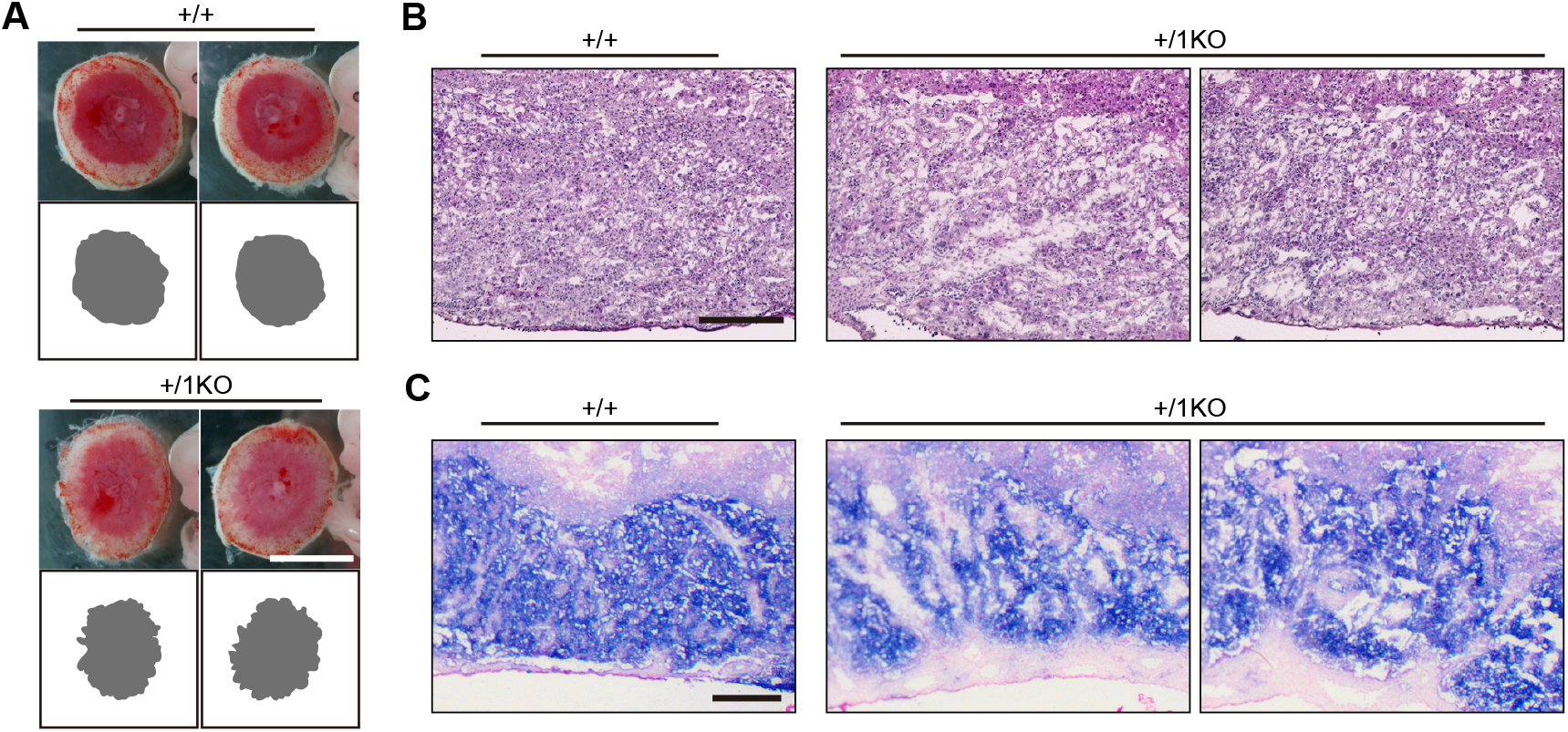
The +/1KO mice exhibit abnormal placental morphologies. (A) Images of the entire placenta of the +/+ (top) and +/1KO (bottom) observed from the fetal side at 13.5 dpc. The labyrinth area is shown filled in gray at the bottom of each image. (B) HE-stained histological sections of the 13.5 dpc placenta. (C) Trophoblast cells in the labyrinth layers at 13.5 dpc were detected by alkaline phosphatase (AP) staining. Scale bars indicate 300 μm.

### Underdevelopment of fetal vasculature in the *Peg10*-ORF1KO labyrinth

Detailed morphological analyses revealed that *Peg10*-ORF1KO placentas exhibit structural abnormalities of the fetal vasculature beginning at its onset of development. As PEG10 expression is restricted to trophoblast cells in the labyrinth and absent from fetal endothelial cells (14), the defects observed in mutants are likely attributable to abnormal trophoblast development. In the placental labyrinth, fetal vasculature consists of endothelial capillaries surrounded by three trophoblast layers: two syncytiotrophoblast layers (SynT-I and -II) and a mononuclear sinusoidal trophoblast giant cell (s-TGC) layer (19, 20). Immunofluorescence for CD71, a SynT-I marker, revealed that mutant placentas displayed a coarser and more simplified arrangement as early as 10.5 dpc, in contrast to the fine, complex, wavy structure of wild-type SynT-I layers (Fig. 4A). Consistent with this, fetal capillaries detected by CD31 staining were significantly reduced and poorly developed in mutans (Fig. 4B).

**Figure 4.**
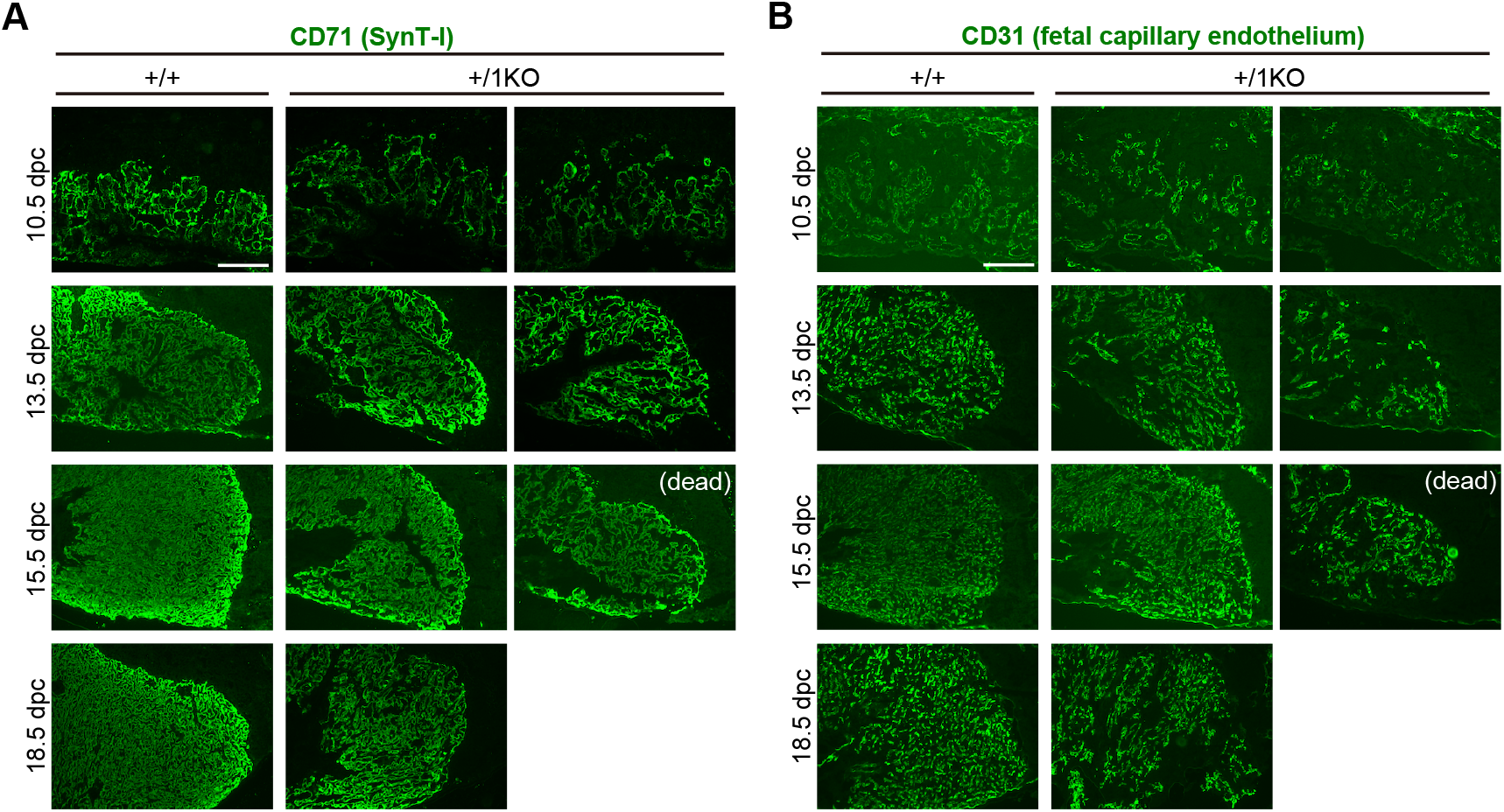
Underdevelopment of the fetal vasculature in the +/1KO placental labyrinth. (A)(B) Immunofluorescence analysis of the placentas at 10.5, 13.5, 15.5 and 18.5 dpc with an anti-CD71, SynT-I marker (A) and CD31, endothelial marker (B) antibodies (green). (dead) refer to the specimens recovered from dead fetus. Scale bars indicate 300 μm.

At 9.5 dpc, when the fetal vasculature is still in a pre-developmental stage, no differences in CD71 or CD31 staining were observed between wild type and mutants; CD71-positive trophoblasts were aligned, and CD31-positive fetal endothelial cells were faintly visible in the chorionic plate (*SI Appendix*, Fig. S5). These findings indicate that vascular abnormalities in mutants emerge concurrently with the onset of fetal vasculature development. Interestingly, the severity of impairment varied among mutant placentas and was most pronounced in those from dead fetuses (Fig. 4A, B). This variability likely accounts for differences in the timing of lethality: fetuses with relatively better-developed labyrinth structures survived longer without significant impacting placental growth, whereas those with severely underdeveloped placentas succumbed earlier. Taken together, these findings suggest that aberrant organization of the fetal vasculature in *Peg10*-ORF1KO placentas profoundly impairs feto-maternal exchange within the labyrinth layer, leading to severe growth retardation and lethality from mid-gestation.

### Abnormal development of trophoblast progenitor cells during labyrinth formation in *Peg10*-ORF1KO

The absence of PEG10-ORF1 protein led to dysregulation of LaTP cells. The variability of placental impairment—some mutant fetuses, retaining near-normal labyrinths and surviving to term (Fig. 4; Table 1)—indicates that ORF1-deficient cells retain partial developmental potential. To identify the underlying defects, we examined characteristics of mutant trophoblasts, including proliferation, apoptosis, and progenitor content. Immunofluorescence using the proliferation marker Ki67, the mitotic marker phospho-histone H3 (Ser10), and the apoptotic marker cleaved caspase-3 showed no clear evidence of altered cell-cycle regulation or apoptosis in mutants (*SI Appendix*, Fig. S6). By contrast, CD326-positive LaTP cells—multipotent progenitors capable of differentiating into SynT-I, SynT-II, and s-TGC (21, 22)—were significantly reduced in mutants from 10.5 dpc, despite appearing normal at 9.5 dpc (Fig. 5). This finding indicates that PEG10-ORF1 positively regulates the onset of LaTP self-renewal and differentiation.

**Figure 5.**
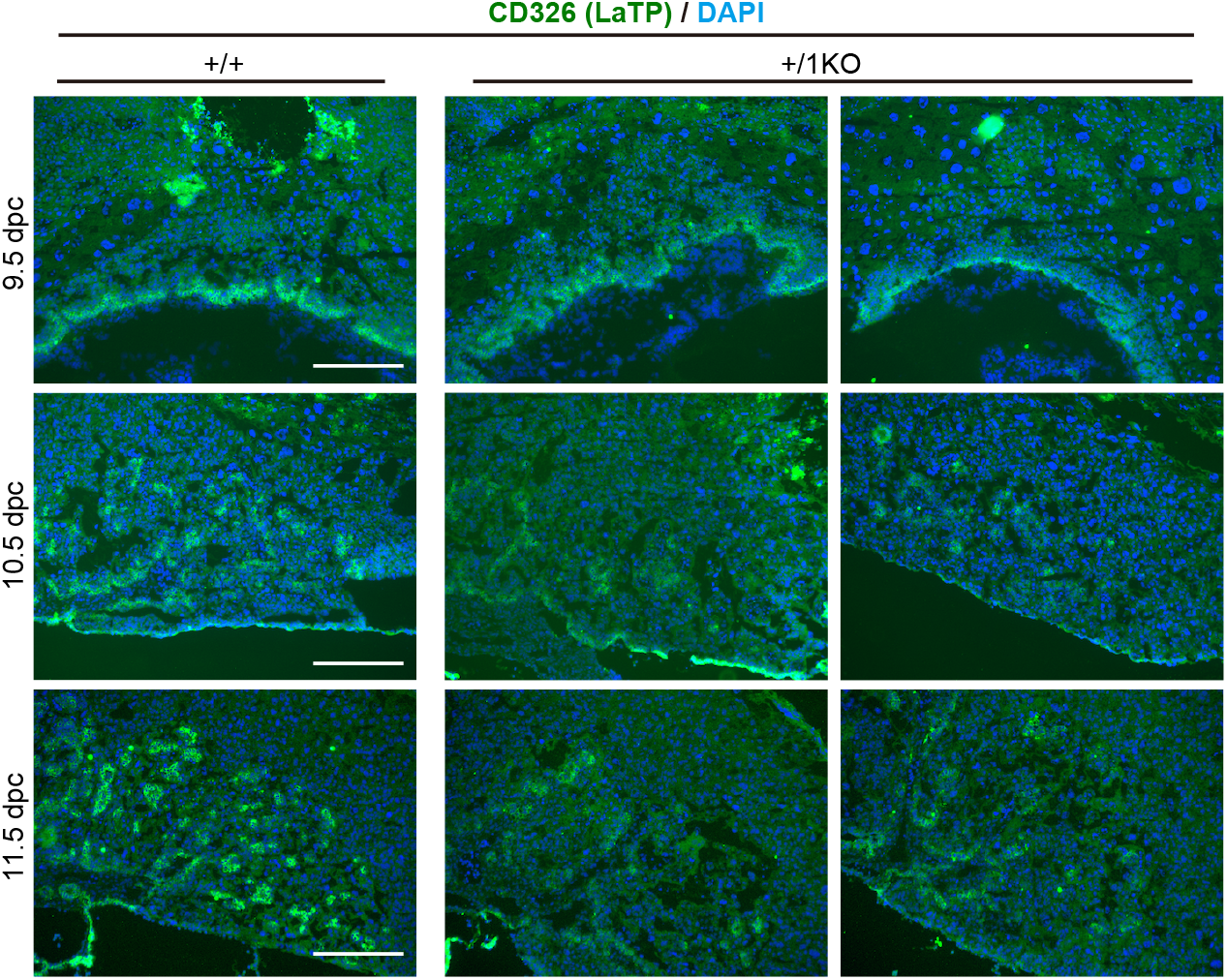
CD326-positive labyrinth trophoblast progenitor (LaTP) cells are obviously decreased in the 1/KO labyrinth at 11.5 dpc. Immunofluorescence analysis of the 9.5 (top), 10.5 (middle) and 11.5 (bottom) dpc placentas with an anti-CD326 antibody (LaTP marker, green). Nuclei were stained with DAPI (blue). Scale bars indicate 300 μm.

Furthermore, detailed immunofluorescence staining for both CD71 and CD31 (Fig. 4A, B) revealed coarser vascular branching and conspicuous avascular spaces in the basal labyrinth adjacent to the chorionic plate (central lower region at 10.5 dpc and lower-left region at later stages in Fig. 4A, B). Based on the results of AP staining (Fig. 3C), these spaces appear to be filled with allantois-derived mesodermal tissue invading the labyrinth. As fetal vascular branching depends on the coordinated differentiation of allantois-derived mesodermal and trophoblast cells (23), this phenotype likely reflects an accumulation of undifferentiated mesodermal cells caused by delayed LaTP differentiation into trophoblast lineages. Thus, PEG10-ORF1 acts as a positive regulator of LaTP differentiation. Variation in the timing and extent of LaTP differentiation among mutants likely determines both the number of differentiated trophoblasts and the maturity of the feto-maternal interface, thereby directly influencing the severity of placental defects and the timing of lethality.

## Discussion

### Importance of the multiple stage-specific functions of *Peg10*-derived proteins and peptides during placentation in eutherians

In this study, we demonstrated that disruption of the PEG10-ORF1 protein leads to mid-gestational lethality accompanied by severe fetal growth retardation, most likely due to delayed differentiation of LaTP cells. This delay resulted in extensive structural abnormalities of the labyrinth and impaired feto-maternal circulation. These findings establish that the ORF1 protein, translated through canonical ribosomal decoding of *Peg10* mRNA, has an essential and distinct role in sustaining placental development and fetal survival during mid-gestation.

Interestingly, ORF1 deficiency also caused a ∼1.7-fold increase in ORF1/2 protein levels (Fig. 1B), despite unchanged *Peg10* transcript abundance (Fig. 2C). This suggests that −1 ribosomal frameshifting not only enables co-translation of ORF1 and ORF1/2 from a single transcript but also fine-tunes their relative expression. Importantly, paternal duplication of proximal chromosome 6 (PatDp(prox6)), which includes the *Peg10* locus, is predicted to double both ORF1 and ORF1/2 protein levels but still yields viable, phenotypically normal mice (24, 25). This supports the conclusion that the phenotype of *Peg10*-ORF1 KO mutants arises from loss of ORF1 function rather than increased ORF1/2 expression. Nonetheless, to fully exclude potential effects of ORF1/2 overexpression, we are pursuing *in vivo* overexpression studies.

Through our series of *Peg10* mutant studies, it is particularly noteworthy that although each mutant exhibits a distinct phenotype, all share the common outcome of severe placental insufficiency leading to high mortality. First, conventional *Peg10* KO mice, which lack all *Peg10*-derived proteins, peptides, and protease activity, show an almost complete absence of the labyrinth and spongiotrophoblast layers, resulting in embryonic death before 10.5 dpc (7). Second, *Peg10*-ASG mutant mice, in which the catalytic activity responsible for self-cleavage is abolished, display progressive fetal and placental growth retardation. Approximately half of these mutants also develop severe inflammation in the fetal vasculature, culminating in perinatal lethality (14). Finally, in the present study, *Peg10*-ORF1KO mutants developed dysplasia of the labyrinthine microstructure composed of fetal capillaries, resulting in mid-gestational lethality with over 80% mortality by 18.5 dpc. Together, these findings clearly demonstrate that *Peg10* performs multiple indispensable functions during placentation in a stage-specific manner (Fig. 6). In early development, the PEG10-ORF1/2 fusion protein most likely directs trophoblast differentiation required for initial labyrinth and spongiotrophoblast formation, as this process remains intact in ORF1KO mutants. During mid-to late gestation, the ORF1 protein contributes to the construction of the feto-maternal interface, enabling efficient exchange of nutrients, waste, and O_2_/CO_2_ between fetal and maternal blood. At late gestation, the *Peg10*-encoded protease protects the fetal vasculature from deleterious insults, thereby ensuring continued circulation. Notably, whereas underdevelopment of the feto-maternal interface is evident from the outset in ORF1KO mutants, in protease mutants the structure initially forms but subsequently collapses, further underscoring the distinct and nonredundant roles of ORF1 and the protease. Future investigations will be required to elucidate the precise molecular mechanisms through which each *Peg10*-derived protein or peptide contributes to placental biology.

**Figure 6.**
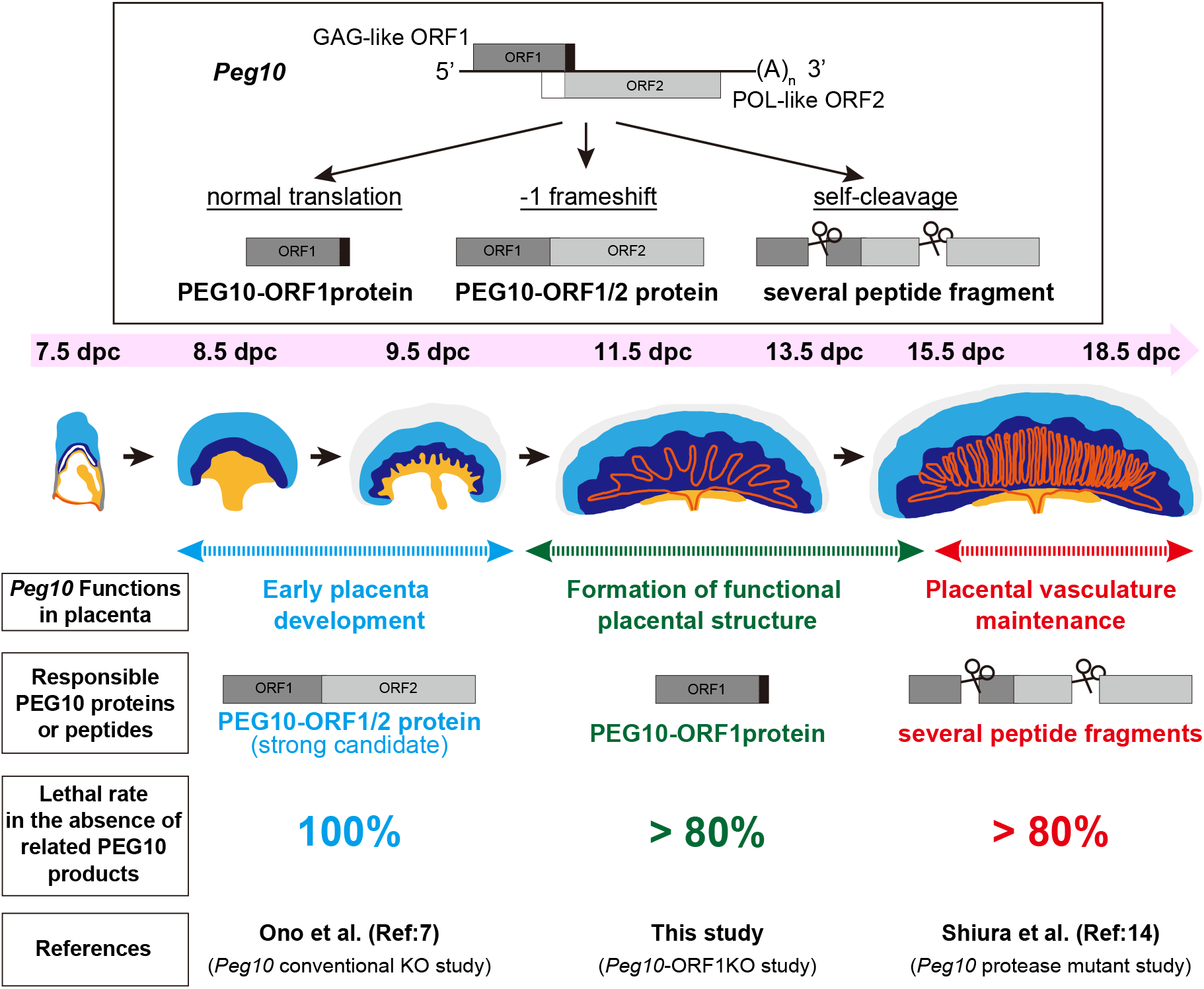
Placental development is governed by the multiple stage-specific functions of Peg10-producing proteins and peptides. The multiple PEG10 proteins and peptides production system originated from an ancestral retroelement (metavirus) enables placental function to be developed and maintained, ensuring normal fetal growth and delivery.

Taken together, our results reveal that *PEG10* has acquired a diverse array of functions essential for placental development by exploiting a multi-protein and -peptide production system—comprising ribosomal frameshifting and protease activity—originally derived from an ancestral retroelement (metavirus) Gag and Pol during molecular evolution. Although it was once hypothesized that the highly invasive hemochorial type evolved secondarily from a non-invasive epitheliochorial form; however, recent phylogenetic evidence suggests the opposite scenario: the ancestral condition was hemochorial (4), or hemochorial or endotheliochorial (26), from which the epitheliochorial type secondarily evolved. A recent ancestral transcriptome reconstruction analysis of uterine endometrial tissue from 23 diverse amniote species also supports the view that the hemochorial type is ancestral (5). Thus, the mouse placenta, with its hemochorial organization, provides an excellent model for understanding the developmental and evolutionary foundations of eutherian placentation. From this perspective, the discovery that *PEG10* exerts multiple stage-specific and essential functions underscores its pivotal role in the origin of eutherian viviparity. Its metaviral origin further highlights how domesticated viral genes were not merely tolerated but harnessed, becoming indispensable drivers of complex organogenesis. The case of *PEG10* exemplifies how the exaptation of viral sequences profoundly shaped the evolution of the mammalian placenta.

## Materials and Methods

### Mice

All animal experiments were approved by the Animal Ethics Committee of the University of Yamanashi. Mice were maintained under specific pathogen-free conditions on a 14:10 h light–dark cycle and were provided free access to a standard chow diet and water.

### Production of *Peg10*-sgRNA

To generate *Peg10*-sgRNA, the template DNA for *in vitro* transcription was synthesized by PCR amplification using the following oligonucleotides: sgRNA-F: GCGGCCTCTAATACGACTCACTATAGGG-GTTCGCCGCCGGGAAACTCCC-GTTTTAGAGCTAGAAATAGCAAGTTAAAATAAGGCTAGTCCGTTATCAACTTGAAAAAGTGGCACCGAGTCGGTGCTTTTTT sgRNA-R: AAAAAAGCACCGACTCGGTGCCACTTTTTCAAGTTGATAACGGACTAGCCTTATTTTAACTTGCTATTTCTAGCTCTAAA.

The resulting T7-sgRNA PCR product was used as the template for *in vitro* transcription with the MEGAshortscript T7 kit (Life Technologies). The synthesized *Peg10*-sgRNA was treated with DNase to remove residual template DNA, purified using the MEGAclear kit (Life Technologies), and eluted in RNase-free water.

### Generation of *Peg10*-ORF1KO mice

B6D2F1 female mice were superovulated, and *in vitro* fertilization was performed using B6D2F1 sperm. The Cas9 protein (Integrated DNA Technologies, IDT; 25 ng/μl), *Peg10*-sgRNA (25 ng/μl), and oligo DNA (50 ng/μl) were co-injected into the cytoplasm of fertilized eggs at the indicated concentrations. The injected embryos were cultured overnight in CZB medium and subsequently transferred into the oviducts of pseudopregnant ICR females. The sequence of the co-injected oligo DNA was as follows: GTGGGGACCTTATTCGTTCTGGCCCTGTCGCTGAAGGTCCCCCTACAGCGGGGCC GGGGA**GATTGCCA**GGCGGCGAATTCTTGGAGGCTTTCGCTGGACACGTGTCGGC GAAATGGCCTCCATTGCCA. Genotyping was performed by restriction fragment length polymorphism (RFLP) analysis of PCR products amplified with the following primers: Forward: GGAAGGTCTCAACCCAGACA, Reverse: GTATCTCACGGTGGTCTCCC. PCR products were digested with BstNI or BciT130I and analyzed by agarose gel electrophoresis. Mutant alleles were identified by the presence of an additional restriction site.

### Real-time quantitative PCR

All assays were performed in triplicate. Gene copy numbers were quantified using the LightCycler 480 system (Roche Diagnostics) with THUNDERBIRD SYBR qPCR Mix (TOYOBO). PCR cycling conditions were as follows: 95 ℃ for 15 s, 65 ℃ for 30 s, and 72 ℃ for 30 s. The primers used were: *Peg10*-F: TTGGTCCCTTACCCCTACCAAC, *Peg10*-R: CCCTTGAGTTAATTCCCAGAGCC, *Actb*-F: AAGTGTGACGTTGACATCCG, *Actb*-R: GATCCACATCTGCTGGAAGG.

### Preparation of frozen sections

Recovered mouse placentas were directly embedded in OCT compound (Sakura Finetek). OCT blocks were cryosectioned at 7 μm thickness (Leica cryostat) and mounted on Superfrost Micro Slides (Matsunami Glass).

### Histological analysis

Frozen sections were fixed in 4% paraformaldehyde for 10 min at room temperature and washed three times with PBS (5 min each). Sections were then subjected to HE staining, immunofluorescence, or endogenous alkaline phosphatase activity detection. For HE staining, sections were stained with HE and mounted with Mount-Quick (Daido-Sangyo). For immunofluorescence, sections were blocked for 60 min at room temperature in blocking buffer (1% bovine serum albumin in PBS containing 0.1% TritonX-100), followed by incubation with primary antibodies in blocking buffer at 4 ℃ overnight.

The primary antibodies used were:

rat monoclonal anti-CD31 (1:100; BD Biosciences, 550274); rabbit polyclonal anti-PEG10-ORF1 (1:100; Cell Signaling Technology, #93531); rat monoclonal anti-CD326 (1:100; BD Biosciences, 552370); rabbit polyclonal anti-Ki67 (1:100; Abcam, ab15580); rabbit polyclonal anti-Phospho-Histone H3 (Ser10) (1:200; Cell Signaling Technology, #9701); rabbit monoclonal anti-cleaved caspase3 (1:400; Cell Signaling Technology, #9664); rat monoclonal anti-CD71 (1:100; SantaCruz, sc-59112).

After three washes with PBS (5 min each), sections were incubated with secondary antibodies in blocking buffer at 4 °C overnight: donkey anti-rat IgG conjugated with Alexa 488 (1:500; Jackson ImmunoResearch, 712-545-153); donkey anti-rabbit IgG conjugated with Cy3 (1:500; Jackson ImmunoResearch, 711-165-152). Sections were washed again in PBS (3 × 5 min) and mounted with VECTASHIELD-Hardset (Vector Laboratories) containing 1 μg/ml DAPI. For endogenous alkaline phosphatase activity detection, after being incubated with 100 mM Tris-HCl (pH 9.5) for 10 min at room temperature, chromogenic detection was performed using NBT-BCIP staining (BCIP/NBT AP substrate [Vector Laboratories]) according to the manufacturer’s instructions. After rinsing with 100 mM Tris-HCl (pH 9.5), sections were counterstained with Nuclear Fast Red (Vector Laboratories). Images were captured with the EVOS M5000 imaging system (Thermo Fisher Scientific).

### Western blot analysis

Placental tissue lysates were separated by SDS–PAGE and transferred onto a PVDF membrane (Bio-Rad) before immunoblotting. Membrane were then probed with rabbit polyclonal anti-PEG10-ORF1 antibody, which detects both PEG10-ORF1 and PEG10-ORF1/2 proteins (1:10,000; produced in-house (14)), followed by incubation with horseradish peroxidase-conjugated goat anti-rat IgG (1:200,000; GE Healthcare, NA935). Signal detection was performed using the ECL system according to the manufacturer’s instructions (Thermo Fisher Scientific).

## Acknowledgments

We especially acknowledge all the members of the T.K laboratory and Advanced Biotechnology Center for the assistance with mouse management and research. This work was supported by the funding program for Grants-in-Aid for Scientific Research (C) (23K05588) to H.S., Grants-in-Aid for Scientific Research (A) (19H00978) to F. I., from Japan Society for the Promotion of Science (JSPS), Nanken-Kyoten (Grant No.2020-40) from Tokyo Medical and Dental University (TMDU) to H.S. and F.I., the Mochida Memorial Foundation for Medical and Pharmaceutical Research to H.S., the NOVARTIS Foundation (Japan) for the Promotion of Science to H. S., and the Paper Output Promotion Project from the University of Yamanashi. Editage reviewed the manuscript prior to submission.

## Author Contributions

H.S., T.K-I. and F.I. designed research; H.S. and M.F. performed research; H.S., M.F., M.O., S.W. and D.I. contributed new reagents/analytic tools; H.S., T.K., T.K.-I. and F.I. analyzed data; and H.S., T.K.-I. and F.I. wrote the paper.

