## Supporting Information for "PEG10-ORF1 programs trophoblast progenitor development for placental labyrinth formation"

**Figures**

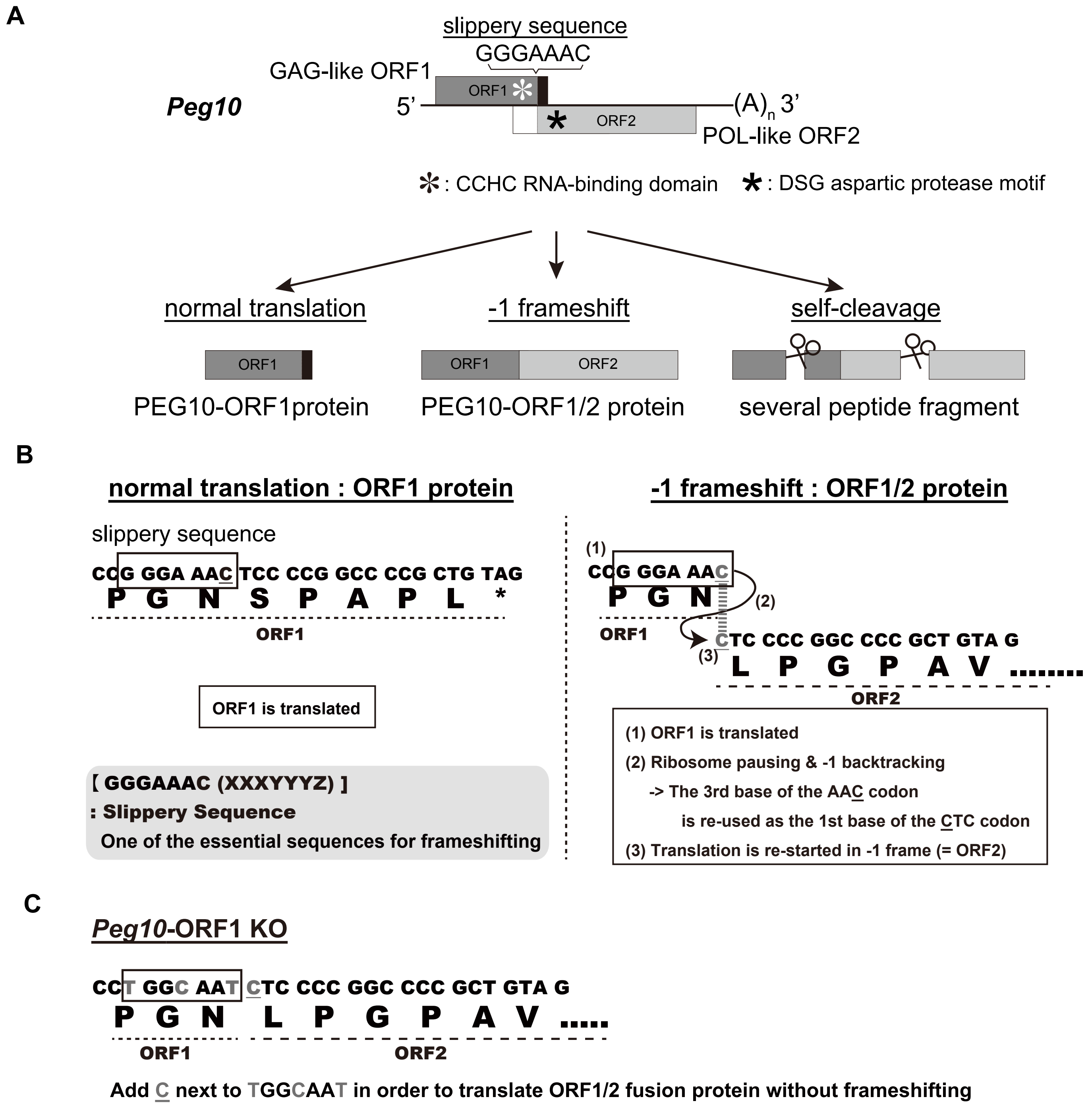


**Fig. S1. The conserved characteristic traits common to retroviruses of PEG10 and *Peg10*-ORF1 KO generation strategy.**

(A) *PEG10* has highly conserved certain characteristic traits common to retroviruses, such as the presence of two ORFs GAG-like ORF1 and POL-like ORF2, a CCHC RNA-binding domain in the GAG-like ORF1 and a DSG viral aspartic protease motif in the POL-like ORF2, and can produce multiple proteins and peptides.

(B) Details of PEG10 frameshifting in mice. *Peg10* produces two different proteins, PEG10-ORF1 by normal translation (left) and the PEG10 ORF1 and ORF2 fusion protein (PEG10-ORF1/2) by using the "−1" translational frameshifting mechanism (right). This example represents the major frameshifting pattern, and other minor frameshifting patterns have also been reported (1). A part of this figure has been reproduced from Shiura et al., 2023 (2) under the CC-BY 4.0 permissions (http://creativecommons.org/licenses/by/4.0/) and include minor modifications and formatting changes from the original figures.

(C) Details of the strategy of *Peg10*-ORF1 KO mice generation. The mutation, GGGAAAC to TGGCAATC, prevents -1 frameshifting and enables translation of only PEG10-ORF1/2 proteins by the normal translation system without changes in amino acid sequence.

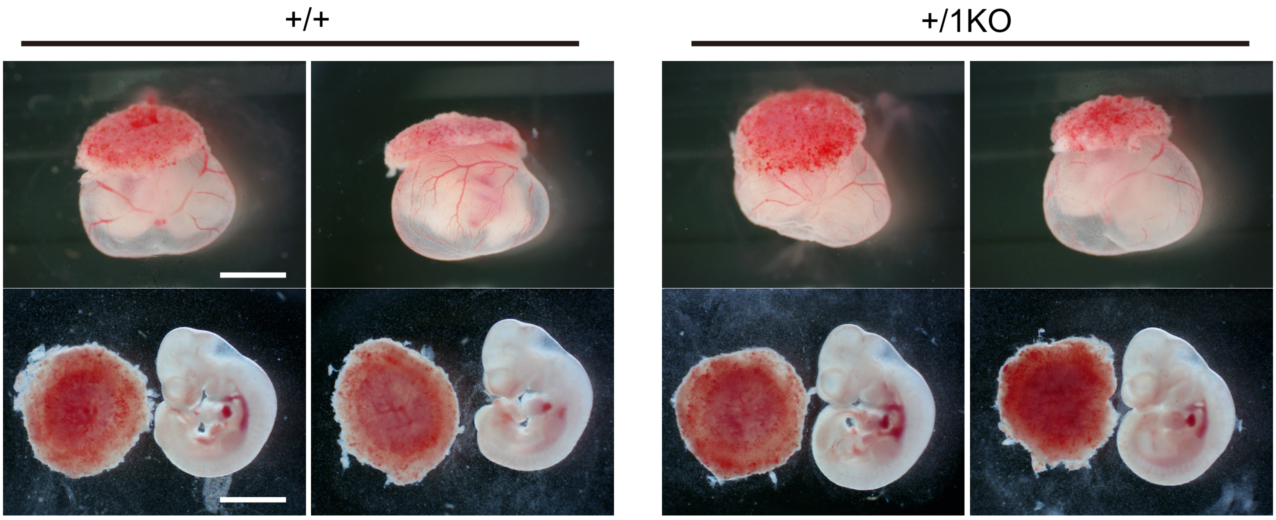


**Fig. S2. Both embryos and placentas of *Peg10*-ORF1KO are apparently normal at 10.5 dpc.**
(Top) +/ + (left) and +/1KO (right) conceptuses and (Bottom) embryos removed from yolk sac along with their placentas at 10.5 dpc. The scale bars indicate 2 mm.

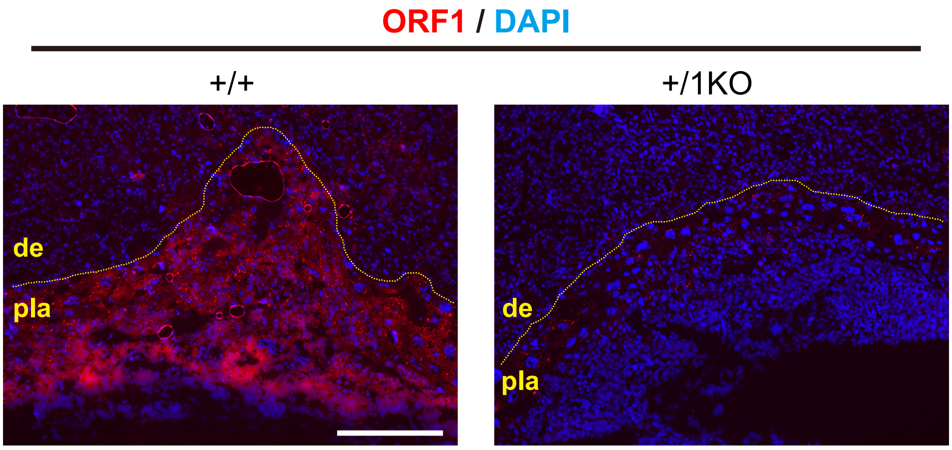


**Fig. S3. PEG10-ORF1 protein expression in 9.5 dpc placenta.**

Immunohistochemical staining with an anti-PEG10-ORF1 antibody. +/ + (left) and +/1KO (right) placenta regions recovered at 9.5 dpc. The dotted lines indicate the borders between the maternal decidua (de) and placenta (pla). The scale bar indicates 300 μm.

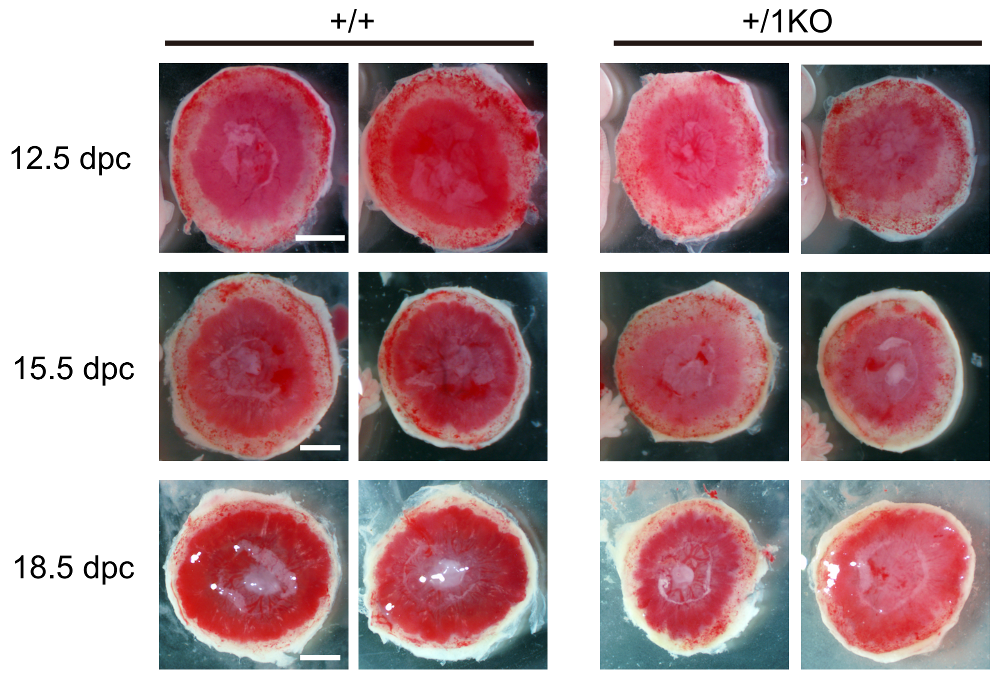


**Fig. S4. Observation of the entire placenta from the fetal side at 12.5, 15.5 and18.5 dpc.**
+/ + (left) and +/1KO (right) placentas recovered at 12.5 (top), 15.5 (middle) and 18.5 dpc (bottom). The scale bars indicate 2 mm.

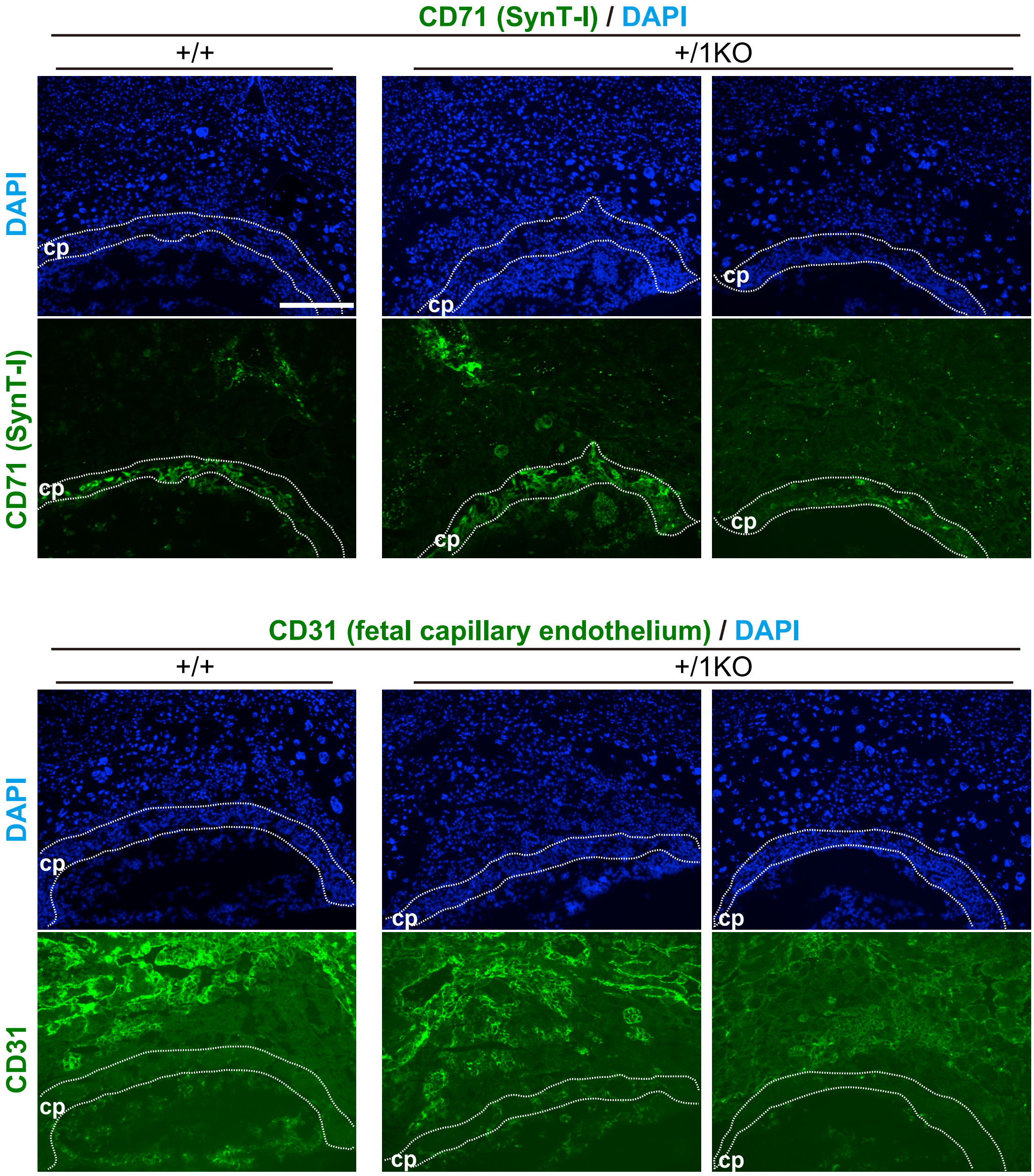


**Fig. S5. CD71 and CD31 expression in 9.5 dpc placenta.**

Immunofluorescence analysis of the 9.5 dpc placenta region with an (top) anti-CD71 (SynT-I marker) and (bottom) CD31 (fetal capillary endothelium) antibodies (green) and nuclear staining with DAPI (blue). Dotted lines indicate the putative chorionic plate (cp). The scale bar indicates 300 μm.

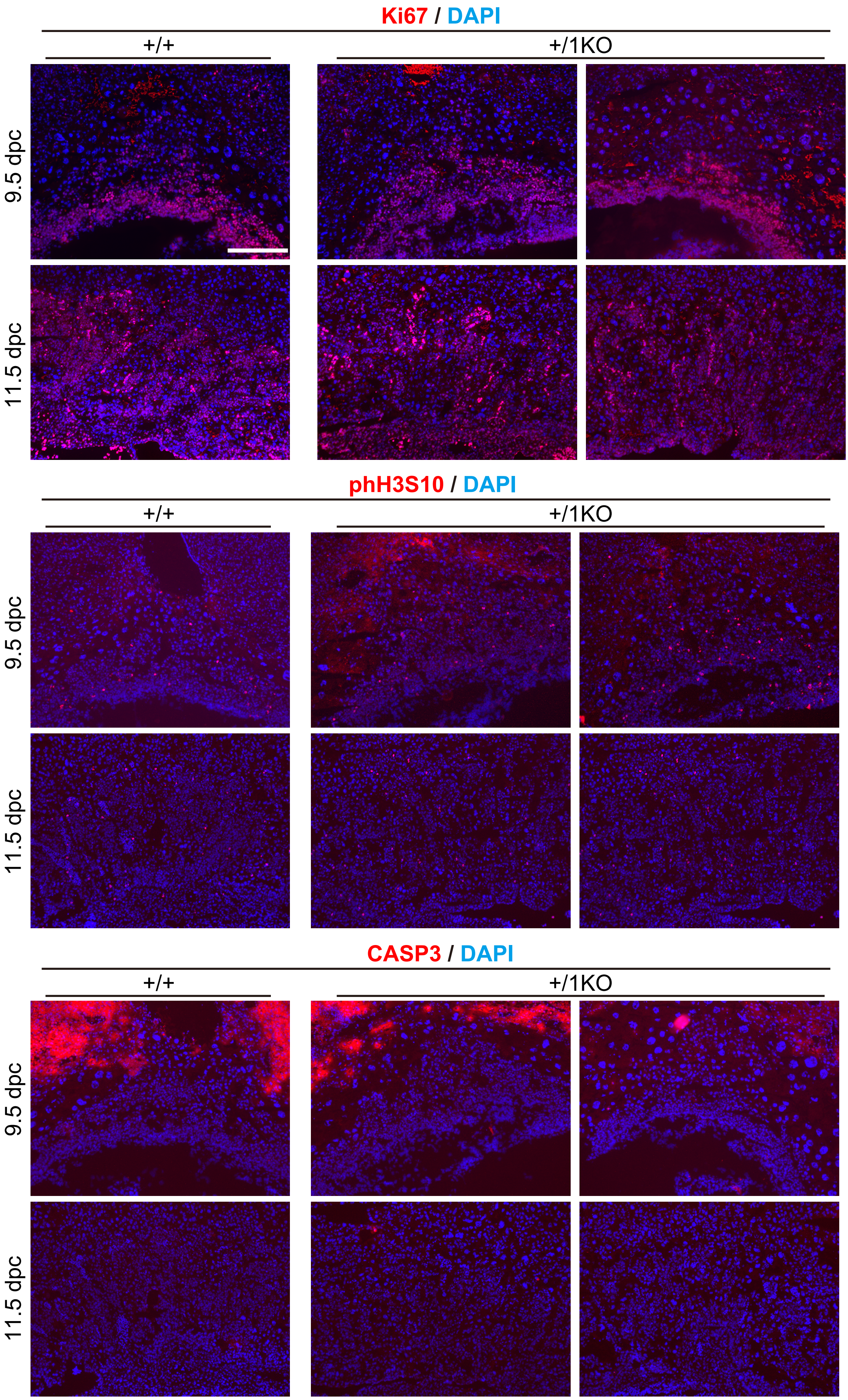


**Fig. S6. No evidence for cell cycle regulation abnormality and apoptosis in the 1KO placenta.** Immunofluorescence analysis of the 9.5 and 11.5 dpc placenta region with an anti-Ki67 (proliferation marker), histone H3 phosphorylated at serine 10 (phH3S10: mitotic marker) and cleaved caspase3 (CASP3: apoptotic marker) antibodies (red) and nuclear staining with DAPI (blue). The scale bar indicates 300 μm.
